# Harmonizing and aligning M/EEG datasets with covariance-based techniques to enhance predictive regression modeling

**DOI:** 10.1101/2023.04.27.538550

**Authors:** Apolline Mellot, Antoine Collas, Pedro L. C. Rodrigues, Denis Engemann, Alexandre Gramfort

## Abstract

Neuroscience studies face challenges in gathering large datasets, which limits the use of machine learning (ML) approaches. One possible solution is to incorporate additional data from large public datasets; however, data collected in different contexts often exhibit systematic differences called dataset shifts. Various factors, *e*.*g*., site, device type, experimental protocol, or social characteristics, can lead to substantial divergence of brain signals that can hinder the success of ML across datasets. In this work, we focus on dataset shifts in recordings of brain activity using MEG and EEG. State-of-the-art predictive approaches on M/EEG signals classically represent the data by covariance matrices. Model-based dataset alignment methods can leverage the geometry of covariance matrices, leading to three steps: recentering, re-scaling, and rotation correction. This work explains theoretically how differences in brain activity, anatomy, or device configuration lead to certain shifts in data covariances. Using controlled simulations, the different alignment methods are evaluated. Their practical relevance is evaluated for brain age prediction on one MEG dataset (Cam-CAN, *n*=646) and two EEG datasets (TUAB, *n*=1385; LEMON, *n*=213). When the target sample included recordings from the same subjects with a different task among the same dataset, paired rotation correction was essential (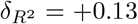 (rest-passive) or +0.17 (rest-smt)). When the target dataset included new subjects and a new task, re-centering led to improved performance (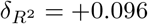 for rest-passive, 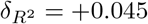 for rest-smt). For generalization to an independent dataset sampled from a different population and recorded with a different device, re-centering was necessary to achieve brain age prediction performance close to within domain prediction performance. This study demonstrates that the generalization of M/EEG-based regression models across datasets can be substantially enhanced by applying domain adaptation procedures that can statistically harmonize diverse datasets.

## 1. Introduction

Magneto- and electroencephalography (M/EEG) are brain recording methods with a high temporal resolution on the order of milliseconds, offering a unique and non-invasive neuroscience method enabling basic research and clinical applications (Hari and Puce, 2017). While quantitative approaches to analyzing M/EEG signals have historically focused on detecting statistical effects, the field has progressively embraced machine learning (ML) approaches whose success is evaluated through predictive modeling. In the context of brain health, classification models are widely used for various applications, *e*.*g*., for epileptic seizure detection (Tzallas et al., 2009), Brain Computer Interface (BCI) (Lotte et al., 2018), or automatic sleep staging (Chambon et al., 2018; Perslev et al., 2021). Even though the regression context has been less explored in the literature, it has been shown to be successful for biomarker learning, *e*.*g*., focusing on brain age as a pretext task (Al Zoubi et al., 2018; Engemann et al., 2020; Sun et al., 2019). In the following, we focus on methods for regression modeling in the particular case of statistical discrepancies between datasets, for example, due to different populations, acquisition devices, or tasks performed during the recording.

Different approaches have been explored to predict cognitive-behavioral or biomedical outcomes from M/EEG data. Methods like Common Spatial Filtering (CSP) (Koles, 1991) or Source Power Comodulation (SPoC) (Dähne et al., 2014) build on top of supervised spatial filtering for dimensionality reduction and unmixing of overlapping, yet physiologically distinct, signal generators. Lately, deep learning based techniques have been the focus of interest as they can learn good feature representation directly from the raw signal, hence potentially simplifying processing pipelines (Roy et al., 2019; Schirrmeister et al., 2017). Independently, an alternative approach has emerged from the BCI community which, like spatial filter methods, summarizes M/EEG data by covariance matrices. But instead of decomposing covariance matrices into filters, this approach uses mathematical tools motivated by the Riemannian geometry of the space of symmetric positive definite (SPD) matrices (Barachant et al., 2012, 2013) to define non-linear feature transformations that facilitate statistical learning with linear models. These techniques perform remarkably well given their simplicity (Congedo et al., 2013; Nguyen et al., 2017; Sabbagh et al., 2020) and are competitive with methods exploiting anatomical information or endto-end deep learning approaches (Engemann et al., 2022). As the field of Riemannian geometry applied to M/EEG is beginning to expand and consolidate, many opportunities remain unexplored. In this work, we focus on investigating the utility of the Riemannian framework for defining dataset-harmonizing transformations.

The recent emergence of large public databases and advances in ML have led to promising prediction models. Yet, these models can be sensitive to shifts in the data distribution and may perform poorly when applied to datasets from other clinical or research contexts. We refer to these gaps between datasets as *dataset* or *domain shifts* (Dockés et al., 2021; Quinonero-Candela et al., 2008). Domain adaptation techniques attempt to deal with these shifts so that a predictive model not only performs well on its source domain but also when applied to another domain (a dataset of a distinct statistical distribution), called the target domain. Many domain adaptation methods exist, ranging from simple approaches (Sun et al., 2017) to more sophisticated models (Damodaran et al., 2018). As we wish to work with M/EEG covariance matrices as basic signal representations for machine learning, we focus on techniques that explicitly use the geometry of SPD matrices to model the statistical distributions of distinct source and target datasets. One first approach proposed in the BCI context is to re-center the distributions to a common point of the SPD space (Li et al., 2021; Yair et al., 2019; Zanini et al., 2018). To get a better alignment of the distributions, (Bleuzé et al., 2021; Maman et al., 2019; Rodrigues et al., 2019) propose to complement this recentering step by adding a re-scale and a rotation correction. These covariance-based methods were initially developed to solve classification problems and are not necessarily applicable to regression without modification. In addition, most of them require labels in the target domain (Bleuzé et al., 2021; Rodrigues et al., 2019) for alignment. We focus on unsupervised alignment methods that can be readily used for regression modeling.

In this paper, we develop a model-based approach for tackling dataset shifts in M/EEG data in which we adapt re-centering, re-scaling, and rotation techniques from previous research on classification (Bleuzé et al., 2021; Maman et al., 2019; Rodrigues et al., 2019) to regression contexts, while assuming that no labeled data is available in the target domain. We build on top of the conceptual framework from (Sabbagh et al., 2020) to investigate how domain shifts can be expressed and handled with an appropriate generative model linking brain activity to both M/EEG measurements and biomedical outcomes. We elucidate how observed dataset shifts can be conceptually decomposed into differences in brain activity and differences in the relationship between the location and orientation of M/EEG signal generators relative to the recording device that reflect the device type, body posture, and individual brain anatomy. With this approach, we establish the connection between particular alignment steps and the parameters of the generative model as well as the physiological and physical shifts they are meant to compensate for. Using statistical simulations, based on the generative model, we then explore different dataset-shift scenarios and investigate the effectiveness of data alignment techniques — combined and in isolation. Through empirical benchmarks on the Cam-CAN MEG dataset (*n*=646) and two EEG datasets (TUAB-normal, *n*=1385; LEMON, *n*=213), we evaluate the practical impact of these alignment techniques for boosting the generalization capacity of regression models across acquisition protocols (resting state vs. audiovisual & motor tasks) and cohorts (clinical EEG versus research & laboratory-grade EEG). We focus on brain age as it is a label easy to collect and valuable as a surrogate biomarker.

The article is organized as follows. In Section 2, we extend the generative model from (Sabbagh et al., 2020) to express and decompose domain shifts into distinct factors, which motivates the three steps we use to compensate for domain shifts: re-centering, re-scaling, and rotation correction. In Sections 3 and 4, we pressure-test these alignment steps using simulations and real-world M/EEG data.

## 2. Methods

To describe dataset shifts that can occur with M/EEG signals, we extend the generative model of M/EEG regression tasks from (Sabbagh et al., 2020) where the prediction target is continuous. A canonical example is brain age prediction (Engemann et al., 2022; Xifra-Porxas et al., 2021). We describe and discuss the parameters of the generative model to understand which mechanisms can explain dataset shifts. Finally, we present various alignment strategies aiming to draw a geometrical analysis of the possible shifts and compensate for them.

### 2.1. Statistical generative model of M/EEG signals

#### Generative model

M/EEG signals **x**(*t*) ∈ ℝ^*P*^ are multivariate time series recorded with *P* sensors at (or above) the surface of the scalp, and that capture the electrical or magnetic activity generated by large-scale neural synchrony. These neurophysiological generators are not directly observable, and here we focus on the situation in which we do not have access to information about the individual brain anatomy, *e*.*g*., when MRI scans are not available. Thus, we use a statistical approach inspired by blind source separation to approximate the signal’s generative mechanism. We model the M/EEG signals as a linear combination of statistical brain generators corrupted by some additive noise. One observation **x**_*i*_(*t*) ∈ ℝ ^*P*^ is written as:

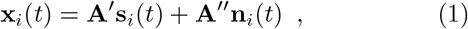

where **s**_*i*_(*t*) ∈ ℝ ^*Q*^ is the underlying signal generating this observation with *Q* ≤ *P*, and 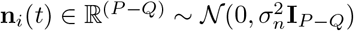 causes a contamination due to noise. We denote **A**^*′*^ = [**a**_1_, …, **a**_*Q*_] ∈ ℝ ^*P×Q*^ the mixing matrix whose columns are the spatial patterns of the neural generators, and **A**” = [**a**_*Q*+1_, …, **a**_*P*_] ∈ ℝ ^*P*×(*P*−*Q*)^ the matrix of the spatial noise patterns. Note that in this model, the noise is not considered independent across sensors but spatially correlated, as is typically the case with environmental or physiological artifacts present in M/EEG data. In this work, we consider datasets for which one observation corresponds to one subject.

This model can be rewritten by combining the generator patterns and the noise in a single invertible matrix **A** = [**a**_1_, …, **a**_*Q*_, **a**_*Q*+1_, …, **a**_*P*_] ∈ ℝ ^*P×P*^ which is the concatenation of **A**^*′*^ and **A**”. The model is thus given by

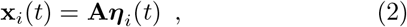

where ***η***_*i*_(*t*) ∈ ℝ ^*P*^ denotes the concatenation of **s**_*i*_(*t*) and **n**_*i*_(*t*). In this model, we assume **A** to not depend on *i* nor *t*. It is also assumed that the statistical generators **s**_*i*_(*t*) = {**s**_*i,j*_(*t*), *j* = 1 … *Q*} are zero-mean, uncorrelated, and independent from the noise. In other words, we assume that the noise generated by artifacts is completely independent of brain activity.

We now consider the covariances **C**_*i*_ of M/EEG signals **X**_*i*_ ∈ ℝ ^*P×T*^ with *T* the number of time samples:

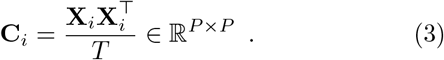

The covariance of M/EEG signals holds the sensors’ variance on its diagonal. In our statistical model and with the previous assumptions, the covariance of the statistical generators is a diagonal matrix whose elements are the variances of each generator 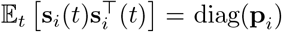 with **p**_*i*_ ∈ ℝ ^*Q*^, also referred to below as “powers”. Thus, we can conveniently summarize the M/EEG covariances as follows:

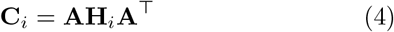

where 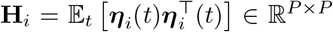 is a block matrix of diag(**p**_*i*_) on the upper *Q×Q* part, and the noise covariance is in the lower (*P* − *Q*) *×* (*P* − *Q*) block. We here assume that 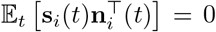, meaning that the matrix **H**_*i*_ is block diagonal.

For regression modeling from M/EEG, it is natural to model the outcome *y*_*i*_ as a linear combination of a function of the generators’ power 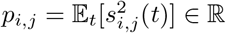:

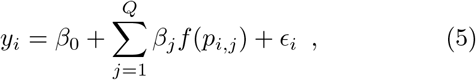

where *β*_*j*_ are regression coefficients, *f* is a known function, and 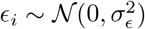 is an additive random perturbation. For example, ageing (*y*) could impact brain activity in distinct brain networks (**s**) to different extents (*β*_*j*_, …, *β*_*Q*_). This could lead for example to a log-decay or log-increase of brain activity per year, hence, motivating a logarithmic function *f* = log, which is a wide-spread function describing the scaling of various facets of brain structure and function (Buzsáki and Mizuseki, 2014) including neural firing rates, axonal diameters, synaptic weights, and, importantly power and frequency scaling.

Replacing the source power with the empirical average of the squared sources, the model is given by:

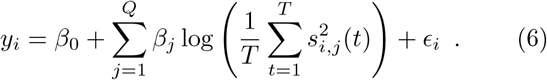

#### Model violations

The assumption that **A** does not depend on the observation (subject) is not valid when working with actual M/EEG data. Each subject has a different head morphology, which results in slight variations in their respective mixing matrices: **A**_*i*_ = **A**+**E**_*i*_ with **E**_*i*_ ∈ ℝ ^*P×P*^. When subscript *i* is omitted below, it will represent the mean observed in the population: **A** represents the average head morphology of the subjects. In our simulations below, we will assume for simplicity that **E**_*i*_ is drawn from 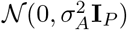.

#### Riemaniann geometry basics

We are working with covariance matrices that belong to the space 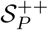 of symmetric positive matrices. These matrices lie in a Riemannian manifold that can be equipped with an appropriate distance (Förstner and Moonen, 2003):

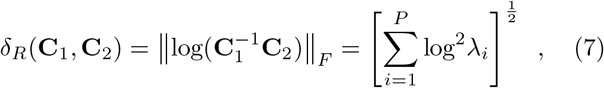

where *λ*_*i*_, *i* = 1, … *P* are the real eigenvalues of 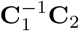. Note that the matrix logarithm of an SPD matrix **C** is computed via its eigenvalue decomposition with the log function applied to its eigenvalues:

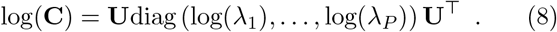

Similarly, the power 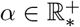 of an SPD matrix is:

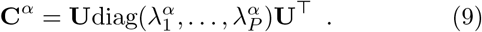

We can then find the geometric mean, or Riemannian mean, of a set of covariances as the minimum of the function:

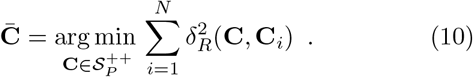

#### Regression method

The approach we focus on in this work involves learning linear models from covariance matrices (Barachant et al., 2012, 2013). Sabbagh *et al*. in (Sabbagh et al., 2019, 2020) show that this Riemann-based model is robust to different preprocessing choices and to model violation. This model also stands out in terms of performance when applied for regression tasks to M/EEG data in various settings.

In this framework, covariances **C**_*i*_ are used as input of the model. The covariance matrices are vectorized to get a feature vector:

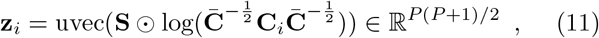

with **S** ∈ ℝ ^*P×P*^ a matrix holding one on the diagonal elements and 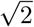 elsewhere, where ⊙ denotes the elementwise matrix product, and uvec the function returning a vector containing the concatenation of the upper triangle values of a matrix. The variables **z**_*i*_ are called tangent vectors, and the matrix **S** preserves the Frobenius norm. We end up with a Euclidean representation of the covariance matrices and can use these vectors as input on classical machine learning models.

### 2.2. Possible data shifts

Each parameter of the model described in equations dataset 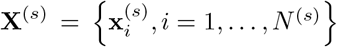 to later predict outcomes on a distinct target dataset 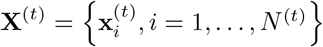 both recorded with the same *P* sensors. The datasets are not necessarily composed of the same number of subjects. Applying our previous notations, we can describe the source dataset as follows:

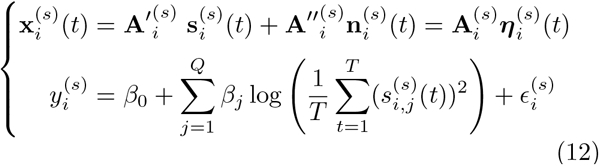

where 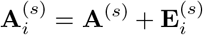 with 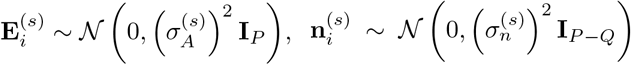 and 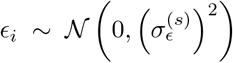. We remind that the statistical generator powers are defined as 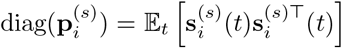. The same equations can be written for the target dataset by replacing the exponent (*s*) by (*t*). We now list physical reasons that could induce differences between source and target datasets and link them to parameter changes of the corresponding generative models (12).

1. If we consider two different populations, the head morphology may vary, and the average subjects would have different mixing matrices: **A**^(*s*)^*≠* **A**^(*t*)^.
2. Having different populations in both datasets would also imply that they will not have the same mixing matrices distribution: 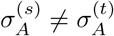
3. When data are recorded with different devices, the recording conditions and noise might not be the same, resulting in different signal-to-noise ratio (SNR): 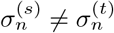.
4. Clinical outcomes *e*.*g*., neuropsychological testing scores can be noisy. This noise could differ from one dataset to another: 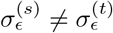.

Because of all those possible causes of variability in the model parameters, machine learning approaches may fail to provide good predictions across datasets. In this work, we focus on shifts that only affect the data, and we assume that the regression coefficients *β*_*j*_ are the same for source and target. In particular, we are interested in understanding changes related to the mixing matrix and the variance of the statistical generators independently of the outcome *y*_*i*_. The variability of these parameters across subjects and datasets affects the observed signals and results in variability in the data distribution. Below, we discuss which statistical methods could help reduce these different gaps or *misalignment* between the data distributions of two different datasets.

### 2.3. Alignment methods

We aim to learn a regression model from one dataset, the source domain, that will perform well on another, the target domain. As we focus on shifts affecting the data distribution, we investigate domain adaptation methods that align the source and the target distributions using geometrical transformations. The methods we chose for understanding and reducing dataset shifts are articulated in three alignment steps: re-centering, equalizing dispersion, and rotation correction. This choice was inspired by transfer learning methods used in Brain-Computer Interfaces (BCI) application and, more specifically, by the Riemannian Procrustes Analysis (RPA) of (Rodrigues et al., 2019). These steps can be used independently, usually by only re-centering the data, or combined. In the following, we detail these alignment functions in a general manner.

#### Step 1: re-centering

The most commonly used method of transfer learning on symmetric positive definite (SPD) matrices is to re-center each dataset in a common reference point on the Riemannian manifold (Li et al., 2021; Zanini et al., 2018). This reference can be chosen as one of the domains’ geometric mean or an arbitrary point on the manifold. Here we propose to re-center each domain to the Identity by whitening them with their respective geometric mean 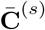 and 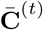:

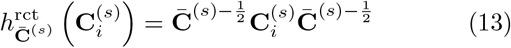

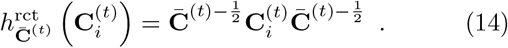

Differently put, re-centering applies separate whitening for source versus target data. This helps avoid errors in the tangent space projection when the average covariance is different for the source and target, *e*.*g*., because the mixing matrices are different, as in Figure 1 (**B**). This is a Riemannian equivalent of the centering step in classical z-scoring.

**Figure 1:**
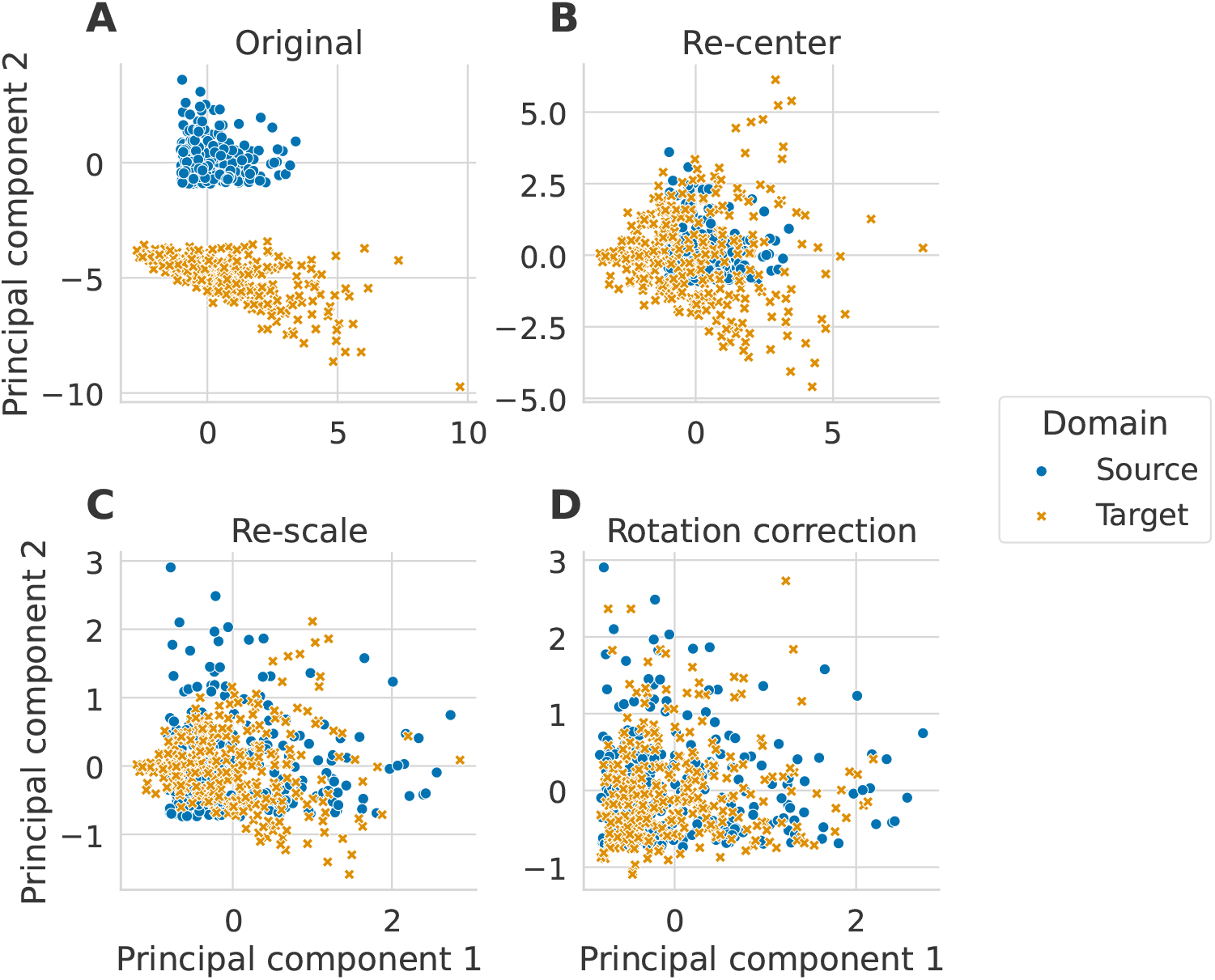
Alignment steps illustrated on simulated data. The three alignment steps are applied to data simulated following the generative model, as detailed in Section 3.1. We set the size of the matrices to *P* = 2 and generated 300 matrices in each domain. Each new step is applied on top of the previous one. The plots correspond to the two first principal components of the tangent vectors. (**A**) The simulated data are plotted on the tangent space before any alignment steps. (**B**) The original simulated data are centered to a common point, (**C**), then their distributions are equalized, and (**D**) finally, a rotation correction is applied.

#### Step 2: equalizing the dispersion

In this second step, the idea is to re-scale the covariances distribution around their mean 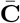 as illustrated in Figure 1(**C**). We first compute the mean dispersion *d* of the covariances as the sum of the square distance between each matrix of the set and their geometric mean 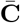 over the number of samples in the dataset:

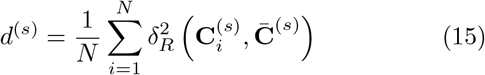

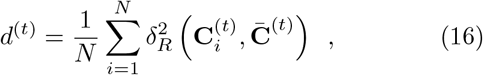

with *δ*_*R*_ the Riemannian distance defined in Equation 7. Then, we re-scale all covariances with 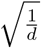 so that the distribution has a dispersion of 1:

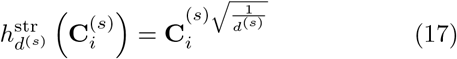

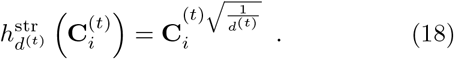

This is a Riemannian equivalent of the re-scaling step in classical z-scoring. Another way to see this preprocessing is to modify the data so that a t-test would not see any difference between the two distributions.

#### Step 3: rotation correction

Until now, we have applied correction measures that process source and target data independently. This is not the case in this third step which implies shared information between source and target datasets. The rotation correction is the most delicate of the three steps. It requires estimating many more parameters than the others, and the source and target feature spaces must be the same size (*P* ^(*s*)^ = *P* ^(*t*)^ = *P*). In the literature, several methods for rotation estimation exist. In the following, we detail two of the methods we selected:

1. The first rotation correction we implemented is inspired by (Maman et al., 2019). The covariances are first vectorized by mapping them in the tangent space (at identity after re-centering). Then we compute the Singular Value Decomposition (SVD) of these tangent vectors 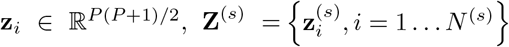 and 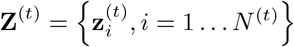:

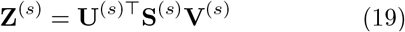

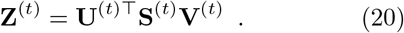

The columns of the **U** ∈ ℝ^*P* (*P* +1)*/*2*×P* (*P* +1)*/*2^ matrices are the left singular vectors ordered from largest to smallest singular values. As the singular vectors obtained with SVD have a sign ambiguity, and since the SVD is done separately for source and target, the resulting source and target SVD coordinates will, therefore, not necessarily match. To orient them identically, we apply a sign correction to the columns of **U**^(*t*)^:

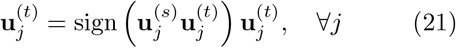

Finally, the **U** matrices are used for rotation correction:

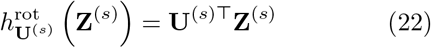

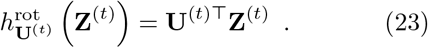

We will refer to this rotation correction method as *unpaired*.

2. The second way to estimate the rotation that we used is inspired by (Bleuzé et al., 2021). In this paper, they consider a classification question and propose to correct the rotation between source and target distributions by matching their respective classes’ mean. This is done by solving the Procrustes problem

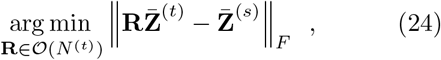

where 𝒪 (*N* ^(*t*)^) is the orthogonal group, 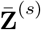 the concatenation of the classes’ mean tangent vector from the source domain, and similarly for 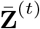 with the available labeled data of the target domain. Then, to correct the rotation, the target tangent vectors are transformed using the solution of the Procrustes problem

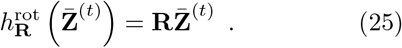

As we wish to be in a regression context without access to target labels, we modified this method by solving the Procrustes problem on all the tangent vectors

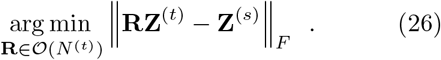

In practice, this solution is found by computing the SVD of the product of the source and target tangent vectors

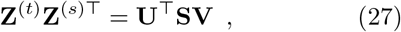

and is **R** = **VU**^⊤^. In this step, we include the information on which source point should be matched to which target point. It means the source and the target dataset should have the same number of observations (*N* ^(*s*)^ = *N* ^(*t*)^ = *N*) and be composed of “matching” observations (for example, the same set of subjects but different tasks/recording conditions/devices). We will refer to this method as *paired*.

This step allows us to align source and target distributions in a shared space. The rotation correction is helpful when the mixing matrices are different between the domains (Fig.1 (**D**)).

## 3. Benchmarks

We conducted a first benchmark with simulated data to evaluate how the alignment steps can compensate for changes in the generative model parameters. Then, we applied the alignment steps to MEG data from the same dataset in two contexts: the same subjects on different tasks and different subjects on different tasks. Then, we evaluated whether the alignment makes it possible to generalize from one M/EEG dataset to another. The regression problem of these benchmarks age prediction. It means we have no noise on the outcome as assumed in Section 2.2. In the following, we detail precisely the different setups.

### 3.1. Simulations

We applied each alignment method in several shift scenarios of the generative model. For each scenario, we varied the magnitude of the shift between the source and the target distributions. We set the dimension of the matrices to *P* = 20 and the number of statistical sources to *Q* = *P*. We considered data with no noise on the signals to be able to easily interpret the performance compared to the one obtained with a perfect model. We did 50 repetitions by generating 300 covariance matrices with 50 random initializations. The target covariances were always a modified version of the source covariances, meaning that one point of the source set corresponds to one point of the target set. This way, it is possible to evaluate the paired rotation correction. Mixing matrices **A** were generated as Gaussian random matrices in ℝ^*P×P*^ from 𝒩 (0, 1). Instead of generating signals **s**, we directly computed their powers **p** as random numbers from a uniform distribution in [0, 1). The same powers were used for both the source and the target sets. We then constructed the covariance matrices 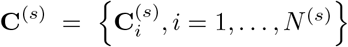 and 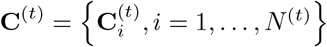, and the outcome to predict as in equations (4) (with **H**_*i*_ = diag(**p**_*i*_) because *P* = *Q*) and (6) (with *ϵ*_*i*_ = 0, ∀*i*).

#### 3.1.1. Simulation scenarios

We detail the changes we introduced for each scenario between source and target distributions. As stated in Section 2.2, we focused on modeling shifts involving changes in the mixing matrix or the variance of the statistical generators. The parameters that are not mentioned were the same for source and target. For the first three scenarios, we aimed to find transformations/shifts between source and target to which each alignment step is robust. Table 2 summarizes the scenarios and the associated parameter changes in source and target.

**Table 1:**
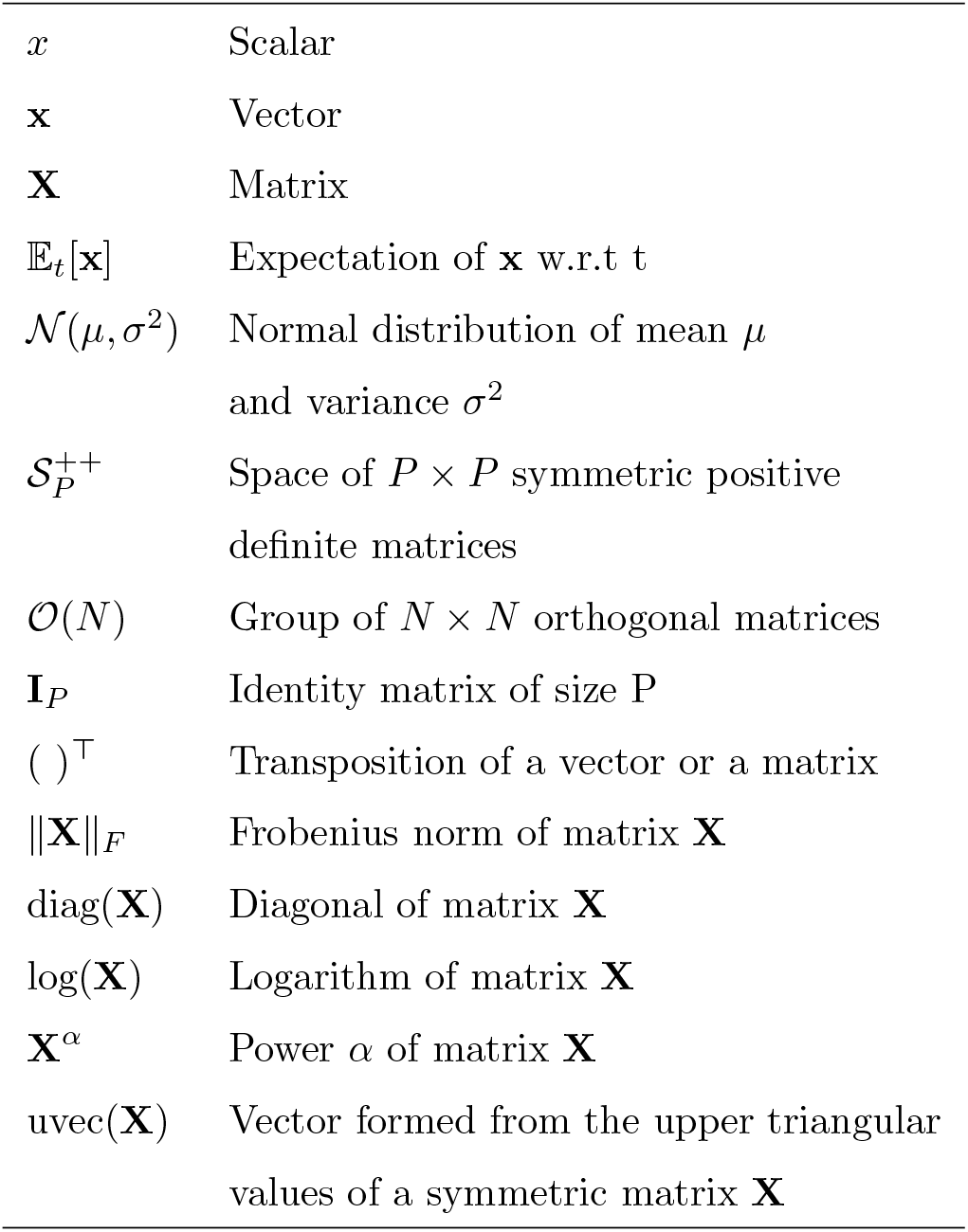
Notations

**Table 2:** Summary of the simulation scenarios.

| Scenarios | Translation | Scale | Translation and rotation | Noise on mixing matrix |
| --- | --- | --- | --- | --- |
| Source | $\mathbf{A}^{(s)}$ fixed | $\mathbf{p}_i^{(s)}$ fixed | $\mathbf{A}^{(s)}$ fixed, | $\mathbf{A}_i^{(s)} = \mathbf{A}^{(s)} + \mathbf{E}_i^{(s)}$<br>with $\mathbf{E}_i^{(s)} \sim \mathcal{N}\left(0, \left(\sigma_A^{(s)}\right)^2 \mathbf{I}_P\right)$<br>$\sigma_A^{(s)} = 10^{-2}$ fixed |
| Target | $\mathbf{A}^{(t)} = (\mathbf{B})^\alpha \mathbf{A}^{(s)}$<br>with $\alpha > 0$ | $\mathbf{p}_i^{(t)} = (\mathbf{p}_i^{(s)})^{\sigma_p}$<br>with $\sigma_p > 0$ | $\mathbf{A}^{(t)} = m\mathbf{A}_t + (1 - m)\mathbf{A}^{(s)}$<br>with $\mathbf{A}_t \neq \mathbf{A}^{(s)}$ fixed<br>and $m \in [0, 1]$ | $\mathbf{A}_i^{(t)} = \mathbf{A}^{(t)} + \mathbf{E}_i^{(t)}$<br>with $\mathbf{E}_i^{(t)} \sim \mathcal{N}\left(0, \left(\sigma_A^{(t)}\right)^2 \mathbf{I}_P\right)$<br>$\sigma_A^{(t)} > 0$ |

##### Translation

In this scenario, we wish to assess how robust the alignment methods are when the source and target mixing matrices are different. Specifically, we built the target mixing matrix as **A**^(*t*)^ = **B**^*α*^**A**^(*s*)^ with 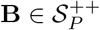 and 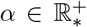. Here, one considered that the mixing matrix perturbation is done by an SPD matrix to decompose the case of **A**^(*t*)^*≠* **A**^(*s*)^ into translation and rotation effects. The benchmark extends the previous simulations from (Sabbagh et al., 2020). More explanations are provided about this decomposition in the **translation and rotation** paragraph. The parameter *α* controls the strength of the perturbation and thus how **A**^(*t*)^ is different from **A**^(*s*)^ (if *α* = 0, **A**^(*t*)^ = **A**^(*s*)^).

##### Scale

We wanted to create a scenario in which the source and target distributions have different dispersions. In this scenario, we constructed the target covariances with an exponent on the powers: 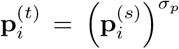 with *σ*_p_ > 0. The parameter *σ*_*p*_ controls how different the dispersions are. This modification was only applied to the data, so the outcome values *y* were unchanged.

##### Translation and rotation

For this scenario, we built the source and target data from completely different mixing matrices and thus generalized the **translation** scenario. To evaluate how alignment methods performed for a growing difference between the source and the target mixing matrices, we defined a parameter *m* such as **A**^(*t*)^ = *m***A**_*t*_ + (1 −*m*)**A**^(*s*)^. **A**_*t*_ was fixed and generated as a random matrix in ℝ^*P×P*^ from 𝒩 (0, 1). In this manner, we created an interpolation between **A**_*t*_ and **A**^(*s*)^ to generate **A**^(*t*)^: if *m* = 0, **A**^(*t*)^ = **A**^(*s*)^ and if *m* = 1, **A**^(*t*)^ = **A**_*t*_. In this scenario, **A**^(*t*)^*≠* **A**^(*s*)^ but we constructed the source and target covariances with the same **H**_*i*_ matrices following equation (4). Thus we can write:

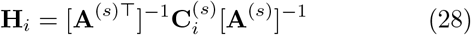

We can replace this **H**_*i*_ expression in the target covariances to get:

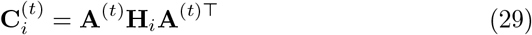

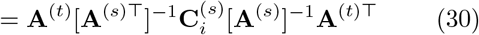

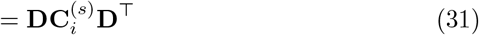

with **D** = **A**^(*t*)^[**A**^(*s*)⊤^]^−1^. The target covariance matrices correspond to the source covariance matrices transformed with the square matrix **D**. A square matrix can be interpreted as a linear transformation: such a matrix can be decomposed into the product of an orthogonal matrix with a positive semi-definite Hermitian matrix (polar decomposition, a.k.a. QR factorization). Thus, we can interpret this scenario as the **translation** scenario (SPD matrix of the polar decomposition) with an additional perturbation by an orthogonal matrix.

##### Noise on mixing matrix

We finally introduced individual noise in the mixing matrix to get a more realistic scenario: 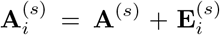 assuming that 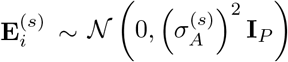 (and similarly for the target mixing matrices). The source data were generated with a fixed noise value 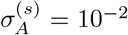, and the tested 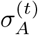 values varied from 10^−3^ to 1. Here, the mean mixing matrices **A**^(*s*)^ and **A**^(*t*)^ were the same. This scenario was inspired by the simulation study of Sabbagh et al. (2020) in which the same level of noise on the mixing matrix was added in the train and the test sets. Here, we explored the situation in which the noise levels in the train (source) and test (target) mixing matrices were different.

#### 3.1.2. Alignment, vectorization, and regression

Once the covariance matrices were generated according to a given scenario, data of both domains were aligned with the methods detailed in Section 2.3. Then, we vectorized the matrices in the tangent space as in (11) with 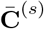 as a reference point for both domains. To avoid numerical issues, we removed low-variance features (see Appendix A for more details). The remaining features were then standardized to get features with zero mean and unit variance. To predict from the standardized vectors in these simulations, for simplicity, we used Ridge regression with its regularization term set to 1. This model was trained on the source data, and predictions were made on the target data. We evaluated these predictions with *R*^2^ scores. Results are presented in Section 4.1 and discussed in Section 5.

### 3.2. M/EEG empirical benchmarks

In the following empirical benchmarks, we focused on one MEG and two EEG datasets for evaluating our alignment methods with real-world data. We first describe these datasets and their preprocessing, then explain how we computed the covariance matrices of the signals. We followed the same preprocessing and processing steps as in (Engemann et al., 2022) for the ‘filterbank-riemann’ pipeline. Finally, we detail the design of each benchmark.

#### 3.2.1. Datasets

##### Cam-CAN MEG data

The Cambridge Center of Aging and Neuroscience (Cam-CAN) dataset (Taylor et al., 2017) consists of MEG recordings from a healthy population covering a wide age range. These data were recorded for each subject during resting state with eyes closed, an audio-visual (passive) task with visual and auditory stimuli presented separately, and a sensorimotor (smt) task with the same stimuli as the previous task combined with a manual response. All data were collected with the same 306-channel VectorView MEG system (Elekta Neuromag, Helsinki) with a sampling rate of 1 kHz.

##### Sample description

We included 646 subjects (319 female, 327 male) with all three recordings. Their age distribution is from 18.5 to 88.9 years with an average of 54.9 *±* 18.4 years and an almost uniform spread over the age range. There was no exclusion of participants. The set of subjects of each benchmark only depends on the availability of recordings for the source and the target tasks and the success of the preprocessing and the feature extraction. Thus some subjects with only two tasks recorded are not included in all benchmarks leading to small variations of the subject sample between benchmarks.

##### Preprocessing

We applied a FIR band-pass filter between 0.1 and 49 Hz to all data. We decimated the signals with a factor of 5 to get a sampling frequency of 200 Hz. To compensate for environmental noise, we performed a temporal signal-space-separation (tSSS) method (Taulu et al., 2005) with a chunk duration of 10 seconds and a correlation threshold of 98%. We only picked channels corresponding to magnetometers (after tSSS signals from magnetometers and gradiometers are mixed and linearly related).

##### TUAB EEG data

The Temple University (TUH) EEG Corpus (Harati et al., 2014) is a large publicly available dataset of clinical EEG recordings. This dataset includes socially and ethnically diverse subjects. In this work, we focus on the Temple University Hospital Abnormal EEG Corpus (TUAB) (Obeid and Picone, 2016), a subset of the TUH EEG Corpus in which recordings were labeled as normal or abnormal by medical experts. Data were collected using several Nicolet EEG devices between 24 and 36 channels and sampled at 500 Hz. The subjects were at rest during the recording.

##### Sample description

We only included healthy subjects with normal EEG in our benchmark. This led to a sample of 1385 subjects (female = 775 and male = 610) with ages between 0 and 95 years (mean = 44.4 years and std = 16.5 years).

##### Preprocessing

Data were band-pass filtered between 0.1 and 49 Hz and resampled to 200 Hz. We selected a subset of 21 channels common to all recording devices used in this dataset. When several recordings were available for one patient, we picked the first to get only one recording per subject.

##### LEMON EEG data

The Leipzig Mind-Brain-Body database provides multimodal data from healthy groups of young and elderly subjects (Babayan et al., 2019). In our benchmark, we only used EEG recordings from this dataset. They were recorded with a 62-channel ActiCAP device and sampled at 2500 Hz. Each subject did two recordings at rest with two conditions: eyes closed and eyes open.

##### Sample description

We included 213 subjects from the LEMON database in our benchmark. No selection criteria were applied, and we kept the data for which the processing and the feature extraction were successful. This led to a cohort with 134 males and 79 females aged from 20 to 77 years. The age distribution of the LEMON presents a peculiarity: it is split into two separate age groups, one with individuals being between 20 and 35 years old and the second between 55 and 77 years old.

##### Preprocessing

A band-pass filter between 0.1 and 49 Hz was applied to the data and resampled to 200 Hz. To keep a maximum of data, recordings with eyes closed and eyes open were pooled before feature extraction.

#### 3.2.2. Data processing and feature extraction

After preprocessing, each filtered recording was segmented in 10 s epochs without overlap. Epochs were then filtered into seven frequency bands as defined in Table 3 as in Engemann et al. (2022). We performed artifact rejection by thresholding extreme peak-to-peak amplitudes on single epochs using the local autoreject method (Jas et al., 2017). Subsequently, we computed covariance matrices from the set of artifact-free epochs with the Oracle Approximating Shrinkage (OAS) estimator (Chen et al., 2010). The ensuing regression pipeline, including all alignment steps, is illustrated in Figure 2.

**Table 3:**
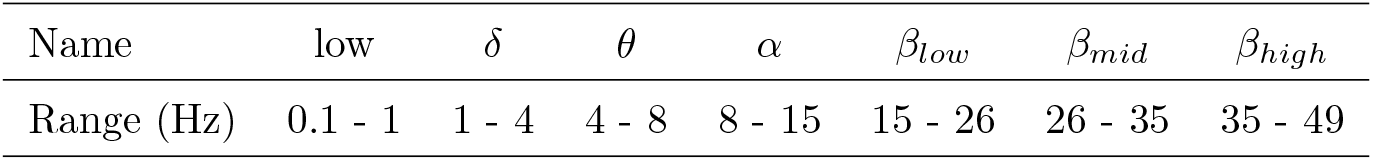
Definition of frequency bands.

**Figure 2:**
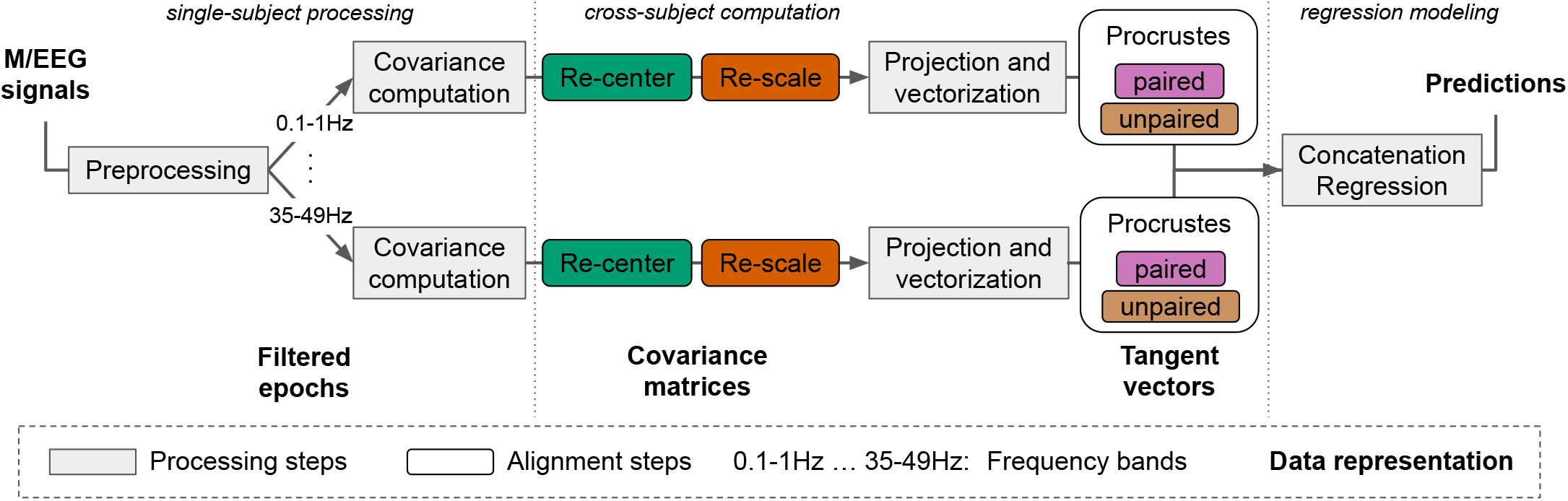
Pipeline for regression modeling with M/EEG with different dataset harmonization steps. For every subject, we summarize the M/EEG recording by the covariance matrix after performing artifact cleaning (Section 3.2.1). The covariances computation, alignments steps, projection to the tangent space, and vectorization steps are done separately for seven frequency bands of Table 3. Alignment steps detailed in Section 2.3 are computed from the covariance distribution across all subjects. The re-center and re-scale steps are performed separately for source and target datasets. The Procrustes steps combine information across source and target datasets. Finally, the seven resulting tangent vectors are concatenated to form one vector per subject used for regression.

For MEG signals, the tSSS method reduces noise by projecting them in a subspace mainly containing the signal, leading to rank-deficient covariance matrices. As a result, it is not possible to correctly apply our alignment methods directly, as rank-deficient covariance matrices are not SPD matrices. To extract valid SPD matrices, we follow the approach from (Sabbagh et al., 2019) and apply Principal Component Analysis to reduce the dimensionality of the covariance matrices, which renders them full rank. We denote the resulting dimension *R*. We use as filters the eigenvectors **W**^(*s*)^ ∈ ℝ^*R×P*^ corresponding to the R highest eigenvalues of the mean source covariance 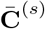. Matrices of both domains are transformed as:

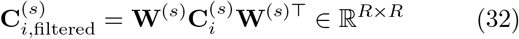

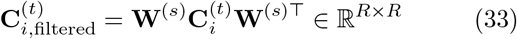

For Cam-CAN data, we set *R* = 65. This spatial filter is applied to each frequency band separately.

#### 3.2.3. Empirical benchmarks

##### Cam-CAN (MEG): same subjects

For this benchmark, we used the experimental tasks of the Cam-CAN dataset for defining the different domains. Here, the source subjects were the same as the target subjects. Only the experimental task (*e*.*g*., audiovisual VS audiovisual + motor) changed from one domain to the other. All subjects included underwent MEG recordings for both the source and the target domain. Therefore, importantly, the mixing matrix and the age distributions were the same for the source and target. As we dealt only with healthy participants, we aimed to minimize the error in age prediction when learning on one task (source domain) and predicting on another (target domain). To estimate standard errors, we did a bootstrap with 100 repetitions.

##### Cam-CAN (MEG): different subjects

In this second benchmark on the Cam-CAN data, we again defined the different domains in terms of the experimental MEG tasks performed by the subjects. Yet, the critical difference with the previous benchmarks is that the source subjects and the target subjects were distinct persons. To implement this analysis, we randomly divided all Cam-CAN subjects into subsets of 80% forming the source subjects, and the left-out 20 % forming the target subjects. A stratification was performed by age decade to maintain similar age distributions between splits. We repeated this split with 100 different random initializations.

##### TUAB & LEMON (EEG): different datasets

In this benchmark, we gauged the performance of alignment methods when the source and target domains are two different datasets. Here, the source domain was composed of data from TUAB, and the target one of data from LEMON. These datasets were not recorded with the same device. However, they had 15 channels in common. We picked the same channels on both datasets to define covariance matrices of the same shape and similar information. The target set was kept fixed, and we implemented a bootstrap procedure on the source subjects to estimate standard deviations.

#### 3.2.4. Alignment, vectorization, and regression

The matrices from both domains were first aligned with the methods described in Section 2.3. We projected the aligned data in the tangent space at 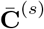 to get tangent vectors. Tangent vectors from all frequency bands were concatenated. Then we applied ridge regression after standardizing (z-scoring) all the features. To select the regularization hyperparameter, we used a generalized (leave-one-out) cross-validation (Golub et al., 1979) on a logarithmically spaced grid of 100 points from 10^−5^ to 10^10^. For quantifying prediction performance, we use the *R*^2^ score.

### 3.3. Software

We processed the M/EEG data with the open-source package MNE-Python (Gramfort et al., 2013), MNE-BIDS (Appelhoff et al., 2019), and the associated MNE-BIDSPipeline (https://mne.tools/mne-bids-pipeline/). The covariance matrices computation and the predictive pipeline were done with the coffeine library (Sabbagh et al., 2020) (https://github.com/coffeine-labs/coffeine). The covariance matrices were manipulated in the alignment methods with the Pyriemann package (Barachant et al., 2022). Results analyses were performed with the ScikitLearn software (Pedregosa et al., 2011).

## 4. Results

We now present the results we obtained from the simulation and M/EEG benchmarks. We first computed the baseline performance of the Riemannian framework without alignment. We then added one alignment step at a time to evaluate its impact on prediction performance. For example, the re-scale method corresponds to re-centering and re-scaling (step 1 + step 2). Likewise, steps 1 and 2 were always performed before rotation correction (Procrustes unpaired and Procrustes paired).

We implemented an element-wise z-scoring to provide comparisons between this widely used method (Apicella et al., 2022; Chen et al., 2021), which is not ideal as it ignores the covariance manifold that can be described and handled by Riemannian geometry, and the presented alignment methods. For this benchmark, we transformed the covariance matrices into correlation matrices by removing the variance of each observation/subject individually. This way, we ended up with symmetric matrices with ones on the diagonal and correlation coefficients between -1 and 1 elsewhere. This transformation corresponds to making the signals zero-mean and with a unit variance before computing the covariance matrices. The correlation matrices were then used as input for the same estimation pipeline as the covariances: projection in the tangent space, vectorization, and regression.

The chance level was represented by a dummy model, which always predicts the mean of the source domain (training data). Performances of all methods were evaluated with the coefficient of determination score or *R*^2^ score, so the higher the score is, the better the model. Scores below 0 indicate predictions performing (arbitrarily) worse than the training mean of the outcome as regressor.

### 4.1. Simulations

We simulated data according to the four shift scenarios detailed in Section 3.1. Figure 3 presents the results for each alignment method and each scenario.

**Figure 3:**
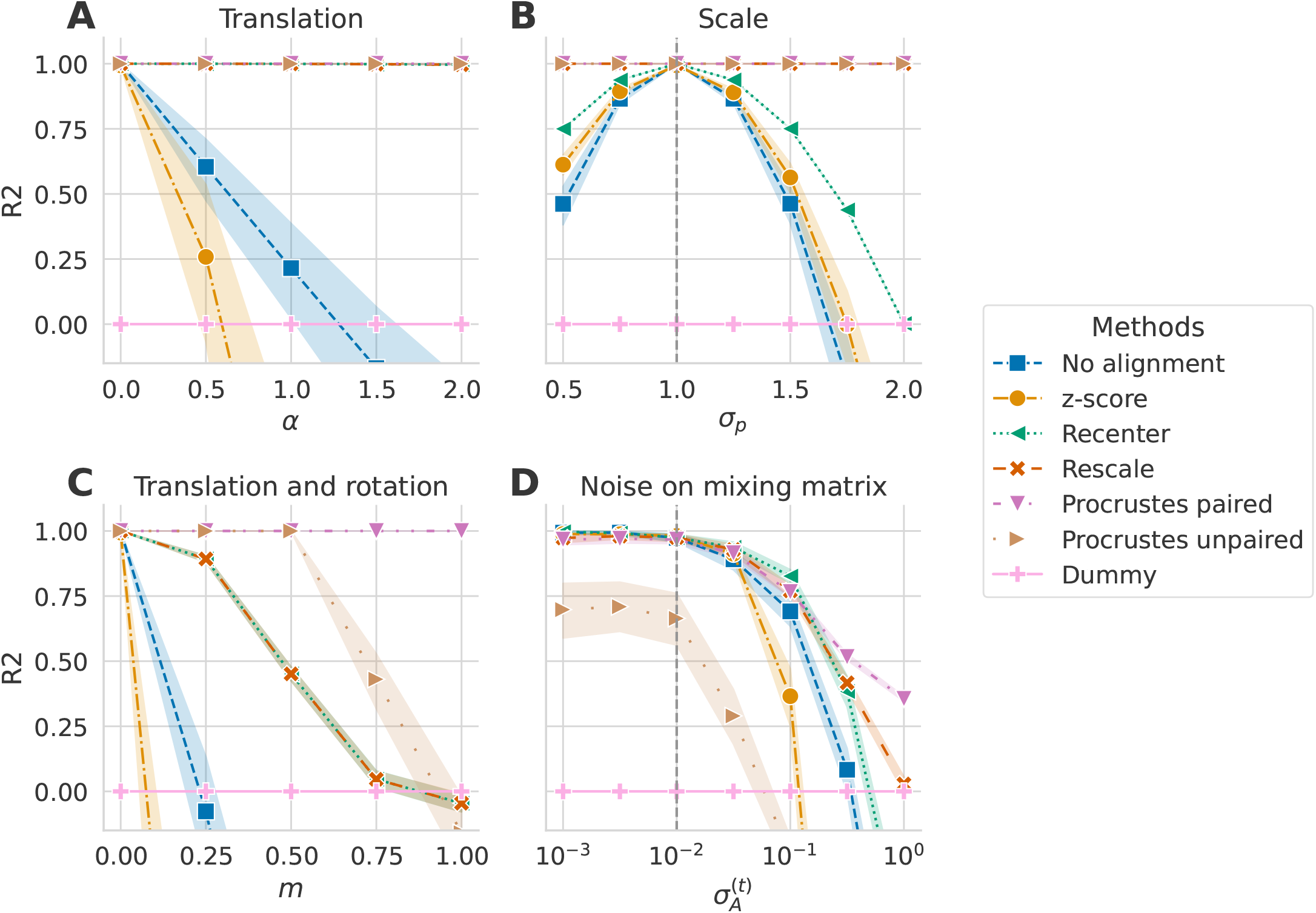
Alignment method comparison across simulated dataset shift scenarios (*R*^2^ score). Alignment methods (indicated by color) were evaluated on four different scenarios with an increasing shift. Error bars show standard deviations of the metric obtained with 50 random repetitions. The dashed vertical gray lines on (**B**) and (**D**) indicate the fixed parameter’s value of the source set. Panel (**A**) displays the performance achieved when the target covariance matrices were created by multiplying the source mixing matrix with an SPD matrix: **A**^(*t*)^ = **B**^*α*^**A**^(*s*)^ with 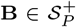. All methods that included re-centering the distributions on the same reference point performed well. (**B**) displays the performance achieved when the dispersion of covariances differs between source and distributions (*σ*_*p*_ ≠ 1). Here, the re-scaling step was essential to align the distributions correctly. (**C**) In this scenario, **A**^(*s*)^ ≠ **A**^(*t*)^, which led to a translation and a rotation of the target set compared to the source set. Re-centering was not insufficient, and a rotation correction was needed to achieve good performance. Interestingly, while Procrustes paired performed well, the unpaired correction broke as the difference between the mixing matrices increased. (**D**) In this scenario, different levels of individual noise were added to the mixing matrices of both domains. For low 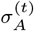 values, all methods except the unpaired rotation correction performed similarly with *R*^2^ scores decreasing slowly. For higher values, the scores dropped, and correcting the rotation with the paired method performed best.

The top left Panel (**A**) illustrates the scores for each alignment method on data generated following the **translation** scenario. The value of *α* controls the shift. As expected, the farther apart the source and target mixing matrices were (higher *α* values), the worse the performance on the unaligned domains method became (in blue). The z-score baseline (light orange) failed even earlier than using Riemannian geometry without alignment. Methods including a re-centering step (green, dark orange, pink, and brown) did not suffer from this shift. This suggests that whitening the source and target distributions by their respective geometric mean mostly compensated for the mixing matrix being perturbed by an SPD matrix. It allowed the regression model to access the log of the powers with little distortion, hence, allowing the linear model to infer the correct function.

Panel (**B**) presents the **scale** scenario in which the log of the target powers were scaled by a parameter *σ*_*p*_ in the signals. When *σ*_*p*_ = 1, the source and target distributions were exactly the same **C**^(*s*)^ = **C**^(*t*)^. In this case, methods including the re-scaling step adjusted for this shift and made accurate predictions, whereas the performance of other methods deteriorated as *σ*_*p*_ increased. Re-centering helped to achieve better predictions compared to no align-ment. The z-score method performed slightly better than not aligning the distribution but was still worse than recentering.

The third Panel (**C**) corresponds to the **translation and rotation** scenario. Here, the target mixing matrix was modified by interpolating between the source mixing matrix **A**^(*s*)^ and another randomly generated matrix **A**_*t*_. The parameter *m* controls where the target mixing matrix is located between these two other matrices, thus how different **A**^(*s*)^ and **A**^(*t*)^ were. The only method reaching perfect predictions, irrespective of the value of *m*, was Procrustes paired. However, the unpaired Procrustes method failed where *m >* 0.5 and even fell behind re-centering. When no rotation correction was applied, re-centering helped to compensate for slight differences between the mixing matrices, but the performance dropped as this difference increased. As expected, a re-centering step and a rotation correction were needed to correct a shift consisting of translation and rotation.

In the last scenario **noise on mixing matrix**, displayed in Panel (**D**), we introduced noise in both source and target mixing matrices to simulate individual differences between subjects. 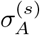 was set at 10^*−*2^, and even,when 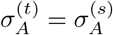 we had 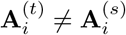. Procrustes unpaired performed worst in this scenario. The unpaired rotation correction was not robust to noise on the mixing matrix. All the other methods performed similarly for low values of 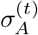. When, all methods deteriorated. The z-score method again showed lower *R*^2^ scores than all other methods. Results suggest that the best solution for this scenario is the paired rotation correction.

The paired rotation correction method performed best in all scenarios but requires the source and target sets to be the same size and have corresponding/paired points. When this is not the case (for datasets with different subjects, for example), re-centering and re-scaling should be the best solution for improving performance. The unpaired rotation estimation seems particularly unstable when the induced shift is too big or when there is noise.

### 4.2. M/EEG data

We now examine the performance of alignment methods with M/EEG data. As with simulated data, the unpaired Procrustes method was not sufficiently robust and led to chance-level performance. We, therefore, do not report it in the following figures.

#### Cam-CAN (MEG): same subjects

We first focused on a regression problem for which there was an individual noise on the mixing matrices, but their distribution was the same in source and target sets because the subjects were the same. It also implies that the age distribution was identical for both domains. The interest of this analysis is to assess what kind of shift is produced when only the task changes and if alignment methods can rectify this. Results for each alignment method on age prediction for three source-target tasks associations are displayed in Figure 4.

**Figure 4:**
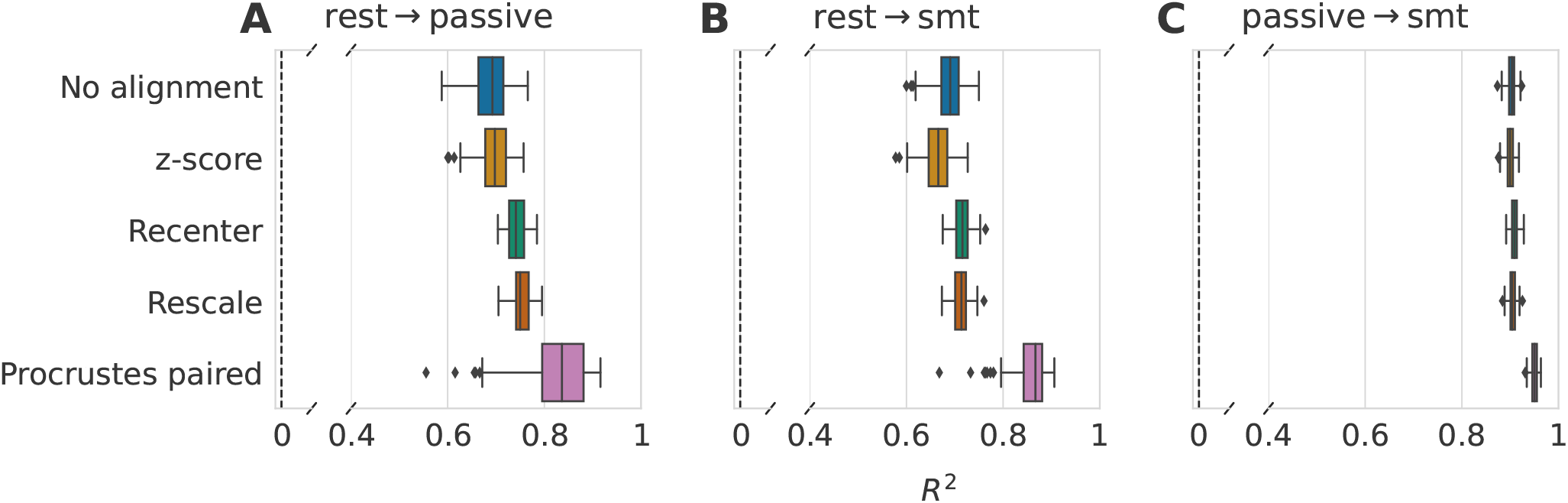
Impact of data alignment on age prediction across different tasks on the same subjects from Cam-CAN dataset (*R*^2^ score). Alignment methods comparison for three different source-target tasks using 100 repeat-bootstrap to select the subjects. Both domains contained the same subjects, only their task was different. Models are depicted along the y-axis, and standard boxplots represent their associated *R*^2^ score. The dashed black lines represent chance-level performance. (**A**) Generalization of age prediction regression model from resting state to the passive task. Re-centering and the paired rotation correction led to clear gains in performance with no obvious benefits for additional re-scaling. (**B**) The regression model was trained on resting-state data, and predictions were made on the recordings of the somatosensory task. Re-centering the data led to slightly improved *R*^2^ scores. Again, the re-scaling step did not lead to further improvements. Correcting the rotation with the paired method contributed to improving 99 splits in comparison to only re-centering. (**C**) Here we used the data from the passive task as the source domain and the somatosensory task as the target domain. Re-centering and re-scaling steps did not affect the prediction performance. The paired rotation correction improved the scores in all splits.

The z-score method led to scores similar to what is obtained without alignment across all three Panels, as expected from the simulation results. For the two first Panels (**A** and **B**), the source domain contained the resting-state recordings, and the target tasks were, respectively, the passive and the somatosensory tasks. The *R*^2^ scores we obtained after these two benchmarks were highly similar. When no alignment was done, the *R*^2^ score was around 0.7. Re-centering the distributions led to a gain in both situations, even though this was more pronounced in Panel (**A**). The re-scaling step had no obvious impact on performance. The paired rotation correction led to improved prediction scores on 83 bootstrap iterations in Panel (**A**) and on 99 iterations in Panel (**B**). In Panel (**C**), the source domain was the passive task, and we made predictions on the somatosensory task, leading to quite different results. The performance reached with no alignment was already very high, with a mean *R*^2^ of 0.9. Re-centering and rescaling gave the same results as not performing any align-ment. Then, the paired rotation correction step induced an apparent gain in performance with a *R*^2^ = 0.95 *±* 0.01. This analysis shows a larger shift between rest and the two other tasks than between the passive and somatosensory tasks. Going from rest to tasks affects the geometric mean of the covariance matrix distributions, but not when going between passive and smt tasks. This suggests that brain activity and, potentially, concomitant physiological artifacts were significantly altered between these tasks. In all three situations, the performance gain obtained with Procrustes paired implies the presence of a rotation of the tangent vector distribution.

#### Cam-CAN (MEG): different subjects

In this second benchmark, we focused on domain-shift differences between MEG tasks in non-overlapping samples of distinct subjects. As a consequence, the distributions of mixing matrices, necessarily, differed for the source and target domains. We performed a cross-validation in which 80% of the subjects were assigned to the source domain and 20% to the target domain. To keep relatively similar age distributions in the train and test splits, we did a stratification on the age decades of the subjects (cf StratifiedShuffleSplit of the Scikit-Learn software). Applying the paired rotation correction in this setup was no longer possible. Thus, it is impossible to analyze whether a rotation exists in the shift. We present the results of each alignment step in Figure 5.

**Figure 5:**
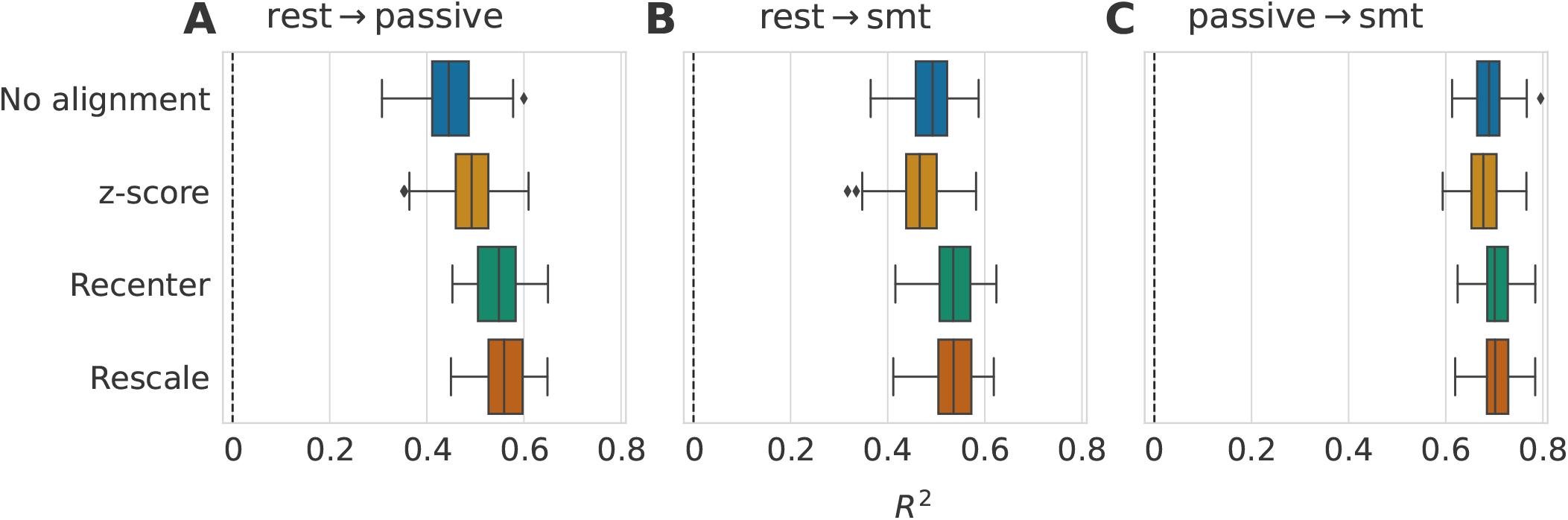
Impact of data alignment on age prediction across different tasks for different subjects from Cam-CAN dataset (*R*^2^ score). Alignment methods comparison for three different source-target tasks using 100 stratified Monte Carlo cross-validation (shuffle split) iterations to determine which subjects form the source and the target sets. We depict the models along the y-axis and represent the *R*^2^ scores with standard boxplots. (**A**) The model was trained on the rest task, and predictions were made on the passive task recordings. When re-centering source and target distributions, prediction performance substantially improved, whereas re-scaling did change performance. (**B**) The target set was composed of recordings from the somatosensory task. The improvement of the re-centering step was smaller but still present. Re-scaling, still, did not lead to obvious improvements. (**C**) In the last Panel, the passive task was the source domain, and the somatosensory task was the target. In this case, aligning was not helpful and led to the same performance as not performing any alignment.

When covariance matrices were not aligned, generalizing from rest to passive tasks led to a *R*^2^ score of 0.55*±*0.05 (**A**), and when the target task was the smt task, we observed an *R*^2^ = 0.54 *±* 0.04 (**B**). The z-score method performed again similarly to the procedure without alignment, with a tendency of slight improvement on Panel (**A**). The re-centering step led to comparable results across generalization scenarios involving resting state and any event-related task (Panels (**A**) and (**B**)). Again, matching the source and target dispersions with re-scaling was not helpful. Finally, all methods showed similar performance when the passive and the smt tasks were the source and the target tasks, respectively (**C**). Our observations for this benchmark match those we made when the subjects were the same for the source and target sets. The *R*^2^ scores reached after alignment in Figure 5 are considerably lower than in Figure 4. Having different subjects in the source and the target domain clearly creates a more difficult-to-reachable shift.

#### TUAB → LEMON (EEG): different subjects

We now consider the resting-state data from two different EEG datasets. The source and target populations are different, as well as the recording devices, but all recordings were done at rest. The source dataset is the larger TUAB dataset (n=1385), and the target dataset is the smaller LEMON dataset (n=213). TUAB also has a broader age range. This way, the regression model will be asked to predict age values that fall within the range observed during model training. We performed a bootstrap with 100 iterations on TUAB data. The results are reported in Figure 6. In addition to the Riemannian approach we focused on in this work (**A**), we were also interested in the impact of the alignment methods on a non-Riemannian model like SPoC (**B**).

**Figure 6:**
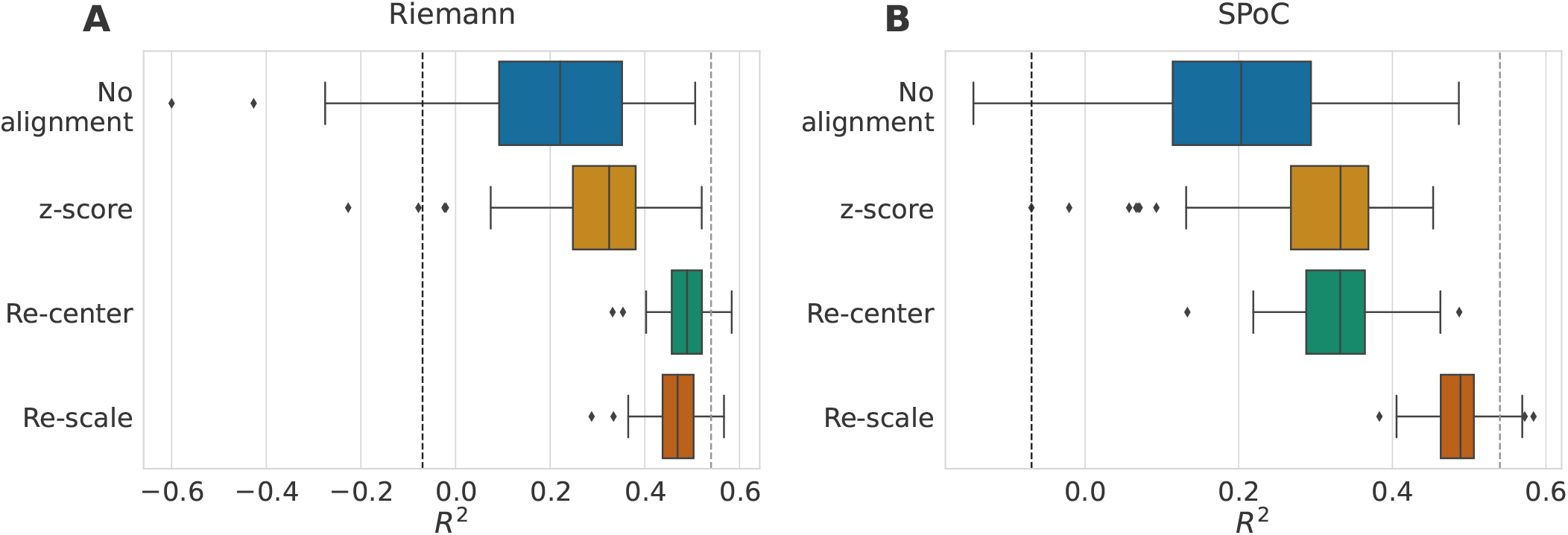
Impact of data alignment on age prediction across different EEG datasets (*R*^2^ score). Data from the TUAB dataset were used as the source domain, and from the LEMON data as the target domain. We compare the alignment methods across 1000 bootstrap iterations on the source data (n=1385). The target set was always the same (n=213). The methods are represented along the y-axis, and we depict their associated *R*^2^ scores with standard boxplots. (**A**) Results of alignment methods combined with the Riemannian approach of Equation 11 (as for all the results we have previously presented). Without alignment, the prediction made on the LEMON data led to *R*^2^ scores far lower than what was reported in (Engemann et al., 2022) (10-fold cross-validation on LEMON data only: 0.54 *±* 0.13 represented by the dashed gray line). When both domains are re-centered to identity, we reached performances similar to when the model is trained on LEMON. Re-scaling did not improve results. (**B**) Results when the regression model follows the SPoC approach. Not aligning led again to poor *R*^2^ scores. Unlike the first panel, the z-score method improved the predictions similarly to re-centering. Re-scaling helped to reach performances closer to those of the Riemannian model trained on LEMON.

Without alignment, the Riemannian model and SPoC led to poor results with mean *R*^2^ scores of around 0.2. On Panel (**A**), the z-score method performed slightly better than no alignment. Re-centering the data drastically improved the age prediction performances with *R*^2^ scores of 0.49 *±* 0.05 with a clearly reduced variance. Adding the re-scaling step on top of re-centering did not bring any improvement in performance, it even slightly degraded it. In (Engemann et al., 2022), the filterbank-riemann pipeline trained on LEMON data only with a 10-fold crossvalidation led *R*^2^ scores of *R*^2^ = 0.54 *±* 0.04. Here, the training sample only consisted of data from the TUAB dataset. The Riemannian re-center step made it possible to reach performance comparable to a model trained within the same dataset. With SPoC (**B**), the z-score method and re-centering led to similar slight improvements in the performance over no alignment. The highest *R*^2^ scores were achieved when the re-scaling step was added to the alignment procedure and almost reached the performance of the Riemannian model trained on LEMON.

Aligning the covariances distribution helped improve prediction performance even with a regression model like SPoC that does not leverage the geometry of the covariance matrices. This observation motivated an examination of how alignment affects the SPoC patterns and the resulting powers. As re-centering is a linear transformation, it is possible to combine it with the SPoC patterns for visualization. Thus this is the alignment method we used for the results displayed in Figure 7. The first two rows of (**A**) illustrate the five first SPoC patterns of unaligned source (TUAB) and target (LEMON) data. Without alignment, the source patterns of the first row were directly applied to the unaligned source and target data, resulting in the log powers represented by the blue dots in the scatterplot (**B**). The target log powers covered a wider range of values than the source log powers and did not match the identity line. We then trained the model on the aligned source data and applied it to the aligned source and target data to get the log powers values represented as orange crosses on the scatterplot (**B**). Re-centering each domain independently resulted in more comparable source and target log powers on average across subjects. To visualize the patterns associated with the aligned log powers and compare them to the unaligned source and target patterns, we displayed on the third row of Figure 7 (**A**) the SPoC patterns of the aligned source data adjusted with the target whitening inverse filter 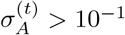. In other words, these adjusted patterns correspond to the SPoC filters applied to the unaligned target data to obtain the target log powers with alignment. The shapes of the adjusted patterns look similar to the source patterns of the first rows with-out any clear transformation in the direction of the target patterns. Even though this analysis was performed in the alpha band, we made the same observations in all other frequency bands.

**Figure 7:**
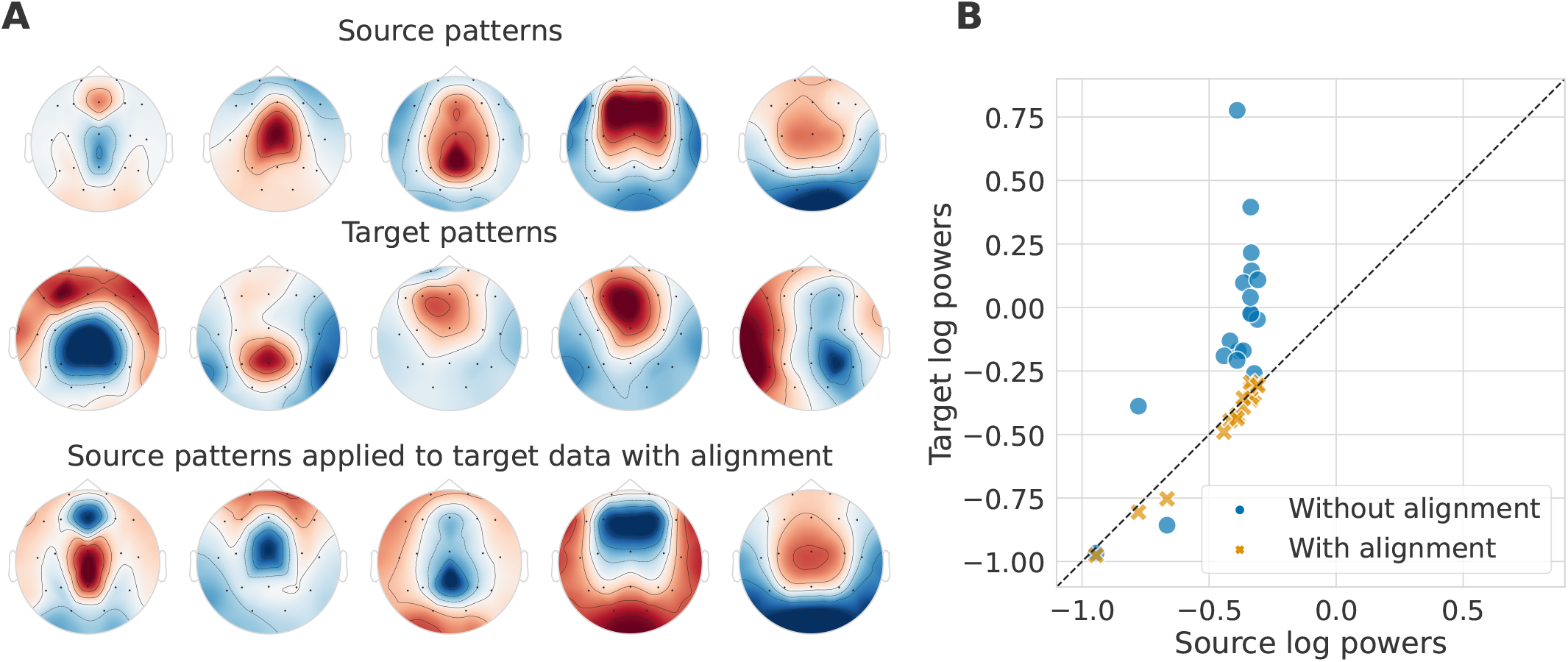
Impact of alignment of different EEG datasets on their SPoC patterns and source powers. TUAB data were used as the source domain, and LEMON data as the target domain. Alignment refers to re-centering the source and the target distribution by whitening them respectively by their geometric mean. To obtain these figures, data were filtered in the alpha band. We included 19 channels (15 commons and 4 with similar locations on the scalp) in both datasets. (**A**) Topographic maps of the five first SPoC source patterns without alignment (first row) and target patterns without alignment (second row). The third row corresponds to the aligned source patterns adjusted with the target whitening inverse filter. These are the patterns applied to unaligned target data to obtain the target powers with alignment. The color map is normalized across each row. (**B**) Scatter plot of the target log powers as a function of the source log powers without and with alignment averaged across subjects. The dashed black line is the identity line. Alignment makes target and source log-powers more comparable.

## 5. Discussion

In this study, we thoroughly explored domain adaptation methods that align M/EEG covariance matrix distributions for regression problems on both simulations and large datasets. We adapted methods from BCI applications (Bleuzé et al., 2021; Maman et al., 2019; Rodrigues et al., 2019) articulated in three alignment steps: re-center the geometric means, equalizing dispersions, and rotation correction. These alignment steps are evaluated in the regression context of generalizing age prediction across different domains. We investigated how dataset shifts can occur by analyzing a statistical generative model of M/EEG data. We presented simulated dataset shift scenarios based on this model for which alignment steps can effectively compensate the shift, plus a noise scenario to get a sense of how the methods would perform with real data. The simulation results showed that Procrustes paired is the most efficient method in all scenarios. It was expected as it includes the three alignment steps and a rotation correction informed by the pairing of source and target subjects. We then designed M/EEG benchmarks with different domain definitions to determine the alignment methods’ efficiency in those various settings. Coherently with the simulation results, Procrustes paired achieved the best performance, but since it cannot be applied in all situations, re-centering is the best option.

We compared the alignment steps leveraging Riemannian geometry with a z-score method that transforms the covariance matrices into correlation matrices. This method systematically performed worse than all the others. Taking into consideration the geometry of the data space is essential. Among the three Riemannian alignment steps, recentering and the paired rotation correction of the source and target distributions help to improve the prediction performance in the M/EEG benchmarks. It was expected in the benchmark where the target population is not the same as the source population, as we showed these steps to compensate for changes in the mixing matrices of the generative model. In the first benchmark on Cam-CAN data, the source and target subjects were the same, but we still observed that re-centering and Procrustes paired led to better scores. On the other hand, equalizing their dispersions did not bring clear gains in performance in any benchmark. In the Cam-CAN benchmarks, the scores reached when the subjects are different in the domains are distinctly lower than with the same subjects: the shift is bigger (lower no alignment baseline) and harder to recover. For the EEG benchmark, we used the TUAB dataset as the source and the LEMON dataset as the target. Here, all recordings were done at rest but with different recording devices and in different populations. Re-centering the distributions in the EEG benchmark exceeded our expectations. Re-centering was sufficient to recover performance close to what is reached when training the Riemannian model on LEMON (Engemann et al., 2022). The re-centering step is simple to implement and has already been very effective in BCI classification to deal with variability between sessions (Barachant et al., 2013) but also between subjects (Zanini et al., 2018). Our results suggest it is also effective in a regression context with variability between populations, tasks, and recording devices.

We extended our evaluation of the impact of alignment methods on different EEG datasets to the SPoC model (Dähne et al., 2014). In this setting, the z-score method and re-centering performed both equally better than no alignment. Interestingly, re-scaling was beneficial and helped to reach performance close to the Riemannian model trained on LEMON. By inspecting the SPoC patterns and the associated log powers, we demonstrated that the observed gain in the performance of re-centering was enabled by more similar log powers between source and target than without alignment. In other words, data alignment adapts the target features to the regression equations fitted on the source data, which explains generalization.

Unfortunately, the two last benchmarks are missing a rotation correction method. As Procrustes paired led to an apparent score increase in the first benchmark, we expect a rotation correction to be beneficial in the other benchmark. Yet, the simulation study showed that the unpaired Procrustes method failed to correctly estimate the rotation when there is noise or when the shift gets too large. The result suggests that this method would likely fail with M/EEG data. The condition of matching subjects between the source and target in Procrustes paired is too restrictive and is not applicable in many settings. The supervised rotation correction methods developed for classification problems (Bleuzé et al., 2021; Maman et al., 2019; Rodrigues et al., 2019) are unsuitable for regression. Further investigations are needed to fill the lack of rotation correction in regression contexts.

Another limitation of this work is that we only performed benchmark that involved source and target covariance matrices formed from the same set of sensors. The dimensionalities of the source and the target data must be equal to apply the predictive model to it. It has been proposed to deal with different dimensionalities of covariance matrices via zero-padding (Rodrigues et al., 2020). However, this method is not applicable if there is no rotation correction afterward, so we could not use it. We also observed that our framework is not robust to sensor permutation in the covariance matrices, even with the same sets of sensors. In our EEG benchmark, we had to select the common channels between the two datasets and sort them to reach acceptable performance even for the re-center step. In addition to leading to a gain in performance, having a proper rotation correction would help to deal with issues related to different numbers or types of sensors in the source and target datasets.

Besides the limitations linked to the rotation correction, further points would deserve future studies. First, we explored unsupervised alignment methods that do not explicitly share any information between the source and the target domains. Comparing our results with supervised methods could allow us to quantify the gain of supervision and to have additional insights into the trade-off between the approaches. A second element to consider is that our goal was to evaluate the alignment methods on a regression problem by minimizing the prediction error. We focused on brain age prediction as age is a label that is easy to collect. But other prediction targets should equally benefit from the methods presented in this work. Importantly, we conducted our benchmarks on healthy participants sampled from the general population. Yet, the biggest impact of our results may be seen when bridging datasets from heterogeneous clinical populations, which remains to be demonstrated.

## Acknowledgements

This work was supported by the grants ANR-20-CHIA-0016, ANR-20-IADJ-0002 and ANR-20-THIA-0013 from Agence nationale de la recherche (ANR).

Numerical computation was enabled by the scientific Python ecosystem: Matplotlib (Hunter, 2007), Scikitlearn (Pedregosa et al., 2011), Numpy (Harris et al., 2020), Scipy (Virtanen et al., 2020) and PyRiemann (Barachant et al., 2022), MNE (Gramfort et al., 2013).

## Author contributions

Conceptualization: A.G., D. E.

Data curation: A.G., A. M., D.E.

Software: A.G., A. M., A. C., D.E., P. R.

Formal analysis: A.G., A.M., A.C., D.E., P. R.

Supervision: A. G., D. E.

Funding acquisition: A. G.

Validation: A.G., D. E.

Investigation: A. M., A. C.

Visualization: A.M.

Methodology: A.G, D.E.

Writing—original draft: A. M.

Writing-review & editing: A. M., A. C., A. G., D. E., P. R.

## Appendix A

### Procrustes unpaired and the VarianceThreshold function

In this appendix, we show that the unpaired Procrustes method should be handled carefully. In particular, we show that the vectorized logarithm maps 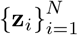, defined in Equation (11), only span a subspace of 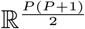 regard-less of the number of covariance matrices **C**_*i*_. Indeed, the rank of 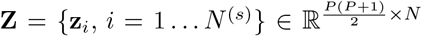 is at most *P*. This implies that, computing the left singular vec-tors 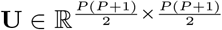, we get 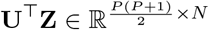 that has at maximum *P* rows with non zero variances. Thus, the other rows must be discarded using, for example, the class `VarianceThreshold` from the scikit-learn library (Pedregosa et al., 2011). Otherwise, numerical issues can be encountered using functions like `StandardScaler` from the scikit-learn library. To prove these assertions, we begin by recalling the mixing model of **C**_*i*_ with no noise in the next hypothesis.

#### Assumption 1

(Mixing model). *We have a set of covari-ance matrices* 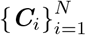 *that respect the following mixing model*

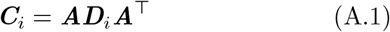

*with* ***A*** ∈ ℝ^*P×P*^ *an invertible mixing matrix and with* ***D***_*i*_ ∈ ℝ^*P×P*^ *diagonal with strictly positive elements*.

This assumption induces that the Riemannian mean 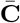 of 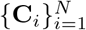 defined in Equation (10) and the associated Riemannian logarithmic mappings 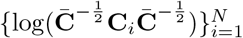 have particular structures. These structures are computed in the following lemma.

#### Lemma 1.

*Knowing* ***A*** *and* 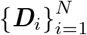, *the Riemannian mean* 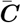 *has a closed form expression which is*

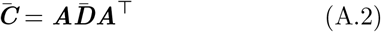

*with* 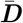 *diagonal with elements* 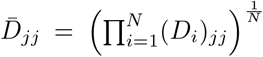 *Furthermore, the Riemannian logarithmic mappings of any* 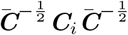 *at identity is*

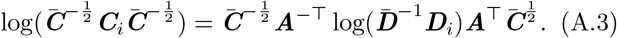

*The log function on the right-hand side of the equation applies the scalar logarithm on the diagonal elements*.

*Proof*. By affine invariance of *δ*_*R*_, we have 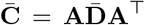 with

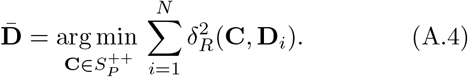

From (Moakher, 2005), 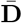 is the unique solution of

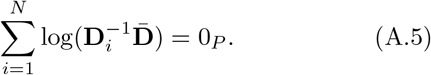

It is readily checked that 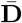 diagonal with elements 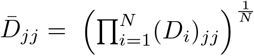 satisfies the Equation A.5. Using these results and the matrix logarithm property log(**EBE**^*−*1^) = **E** log(**B**)**E**^*−*1^ for any **E** ∈ ℝ^*P×P*^ invertible and **B** ∈ ℝ^*P×P*^ such that log(**B**) and log(**EBE**^*−*1^) exist, we have

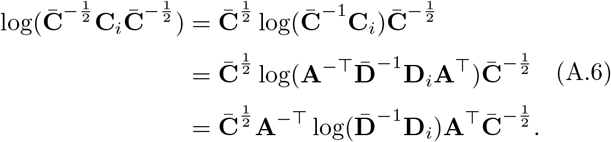

These structures induce that the vectorized logarithm maps 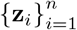 only span a subspace of 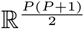.

#### Proposition 1.

*The matrix* 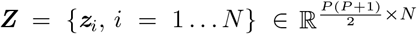 *has a maximum rank of P*.

*Proof*. We begin by defining the full vectorization counterpart of (11)

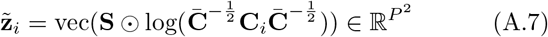

where vec is the operator that concatenates the columns of a given matrix. Then, by denoting **s** = vec(**S**), we get

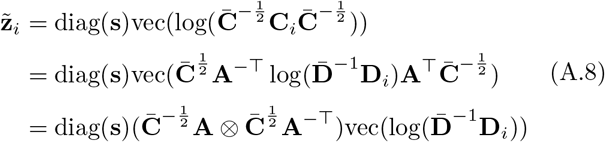

where ⊗ is the Kronecker product. Denoting 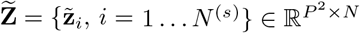, it follows that

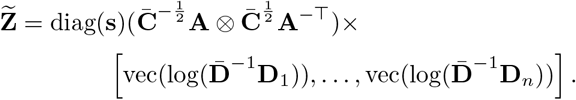

Since rank 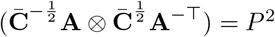, we have that

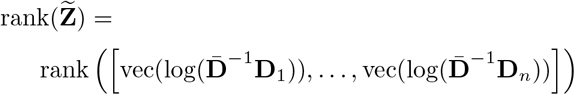

Since log 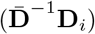 has at most *P* non-zero elements,

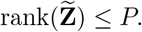

To conclude, the rows of **Z** are included in those of 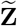, hence

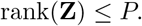

